# Neuropathological validation of the MDS-PSP criteria with PSP and other frontotemporal lobar degeneration

**DOI:** 10.1101/520510

**Authors:** Stefano Gazzina, Gesine Respondek, Yaroslau Compta, Kieren S.J. Allinson, Maria G. Spillantini, Laura Molina-Porcel, Mar Guasp-Verdaguer, Shirin Moftakhar, Stephen G. Reich, Deborah Hall, Irene Litvan, Günter U. Höglinger, Movement Disorder Society-endorsed PSP Study Group, James B. Rowe

## Abstract

**Background:** Progressive supranuclear palsy (PSP) is clinically heterogeneous. Clinical diagnostic criteria were revised in 2017, to increase sensitivity and operationalize the diagnosis of PSP Richardson’s syndrome (PSP-RS) and “variant” syndromes (vPSP).

**Objectives:** To determine the (1) sensitivity and specificity of the 1996 NINDS-SPSP and 2017 MDS-PSP criteria; (2) false positive rates in frontotemporal dementia with frontotemporal lobar degeneration (FTLD); and (3) clinical evolution of variant PSP syndromes (vPSP).

**Methods:** Retrospective multicenter review of 108 neuropathologically-confirmed PSP patients and 81 patients with other forms of FTLD: 38 behavioral variant frontotemporal dementia (bvFTD), 14 non-fluent/agrammatic variant primary progressive aphasia (nfvPPA), and 29 corticobasal degeneration (CBD), Sensitivity and specificity of the MDS-PSP criteria were compared to the NINDS-SPSP criteria at baseline. In a subset of cases, the timing and frequency of clinical features were compared across groups over six years.

**Results:** Sensitivity for recognition of probable and possible PSP pathology was higher by MDS-PSP criteria (72.2-100%) than NINDS-SPSP criteria (48.1-61.1%). Specificity was higher by NINDS-SPSP criteria (97.5-100%) than MDS-PSP criteria (53.1-95.1%). False positives by MDS-PSP criteria were few for bvFTD (10.5-18.4%) but common for CBD and nfvPPA (fulfilling “suggestive of’ PSP). Most vPSP cases developed PSP-RS-like features within six years, including falls and supranuclear gaze palsy, distinguishing frontal presentations of PSP from bvFTD, and speech/language presentations of PSP from nfvPPA.

**Conclusions:** The 2017 MDS-PSP criteria successfully identify PSP, including variant phenotypes. This independent validation of the revised clinical diagnostic criteria strengthens the case for novel therapeutic strategies against PSP to include variant presentations.

## INTRODUCTION

Progressive supranuclear palsy (PSP) was first described as an adult onset, progressive condition characterized by supranuclear vertical ophthalmoplegia, pseudobulbar palsy, axial rigidity, dystonia, and dementia.^1^ Neuropathological examination reveals a characteristic 4-repeat tauopathy most severely affecting subcortical structures such as the globus pallidus, subthalamic nucleus, red nucleus, substantia nigra, periaqueductal grey matter and dentate nucleus.^2^ The presence of tufted astrocytes and the lack of astrocytic plaques allow to distinguish PSP from corticobasal degeneration (CBD).^3^ In 1996 the National Institute of Neurological Disorders and Stroke and the Society for PSP proposed and validated a set of clinical diagnostic criteria (NINDS-SPSP, including slowing of vertical saccades and falls within the first year), that demonstrated good sensitivity (50-83%) and excellent specificity (93-100%) compared to previously published criteria.^4^

However, since the first description of the “classical” PSP phenotype (now generally called Richardson’s syndrome, PSP-RS), it has become clear that many neuropathologically-confirmed PSP cases present with features that were not included, or were considered as exclusionary, in the NINDS-SPSP criteria. Some of these clinically variant PSP syndromes (vPSP) develop features of PSP-RS later in their course, but not all.^5–10^ A recent retrospective multicenter study of 100 pathologic cases found that more than half of PSP cases presented with atypical features, or features not meeting NINDS-SPSP criteria.^11^ Although, pathologic series may include more atypical cases than the range observed in healthcare practice, these results suggested the need for revised criteria that encompassed different presentations of PSP pathology.

Furthermore, “PSP-like” presentations have been reported from other neuropathologies including CBD, frontotemporal dementia, Parkinson disease, motor neuron disease, and cerebrovascular disease, leading to false-positive diagnoses^12^ and underlining the potential for clinical overlap with α-synucleinopathies and frontotemporal lobar degeneration disorders (FTLD), Despite the excellent specificity of the NINDS-SPSP criteria at autopsy, the sensitivity is also limited due to their focus on the PSP-RS phenotype. Several years may elapse before the unequivocal presence of the cardinal features in the additional PSP phenotypes.^5, 6, 13, 14^ vPSP syndromes have been under-recognized^11^, and yet would be relevant to clinical trials of new agents against PSP.

In an attempt to overcome these issues, the International Parkinson and Movement Disorder Society revised the clinical diagnostic criteria for PSP (MDS-PSP).^15^ The new criteria are intended for use both in clinical and research practice. Their aim is to support (i) earlier accurate diagnosis of PSP; (ii) better recognition of vPSP syndromes, and (iii) adding a new “suggestive of’ PSP level of certainty, suitable for early identification of individuals in whom the diagnosis may be confirmed as the disease evolves.^15, 16^

An important feature of the new criteria is the operationalization of vPSP syndromes, including PSP with predominant frontal presentation (PSP-F), PSP with predominant speech/language disorder (PSP-SL), PSP with predominant corticobasal syndrome (PSP-CBS), PSP with predominant ocular motor dysfunction (PSP-OM), progressive gait freezing (PSP-PGF), PSP with predominant parkinsonism (PSP-P), and PSP with predominant postural instability (PSP-PI). The syndromes with prominent parkinsonian and freezing features (i.e. PSP-P and PSP-PGF) have been more extensively recognized and studied than the FTLD-related syndromes (PSP-F, PSP-SL, PSP-CBS). Indeed, it is relevant to determine whether the new criteria for vPSP could be distinguished from their corresponding FTLD-associated syndromes.

The diagnostic value of the clinical syndromes included in the new criteria was determined from neuropathologic cases.^17^ However, independent neuropathological validation is required. This study therefore analyzes an independent multicenter cohort of pathologically-confirmed cases of PSP and other FTLDs, in order to quantify: (1) the sensitivity and specificity of the NINDS-SPSP and MDS-PSP criteria for PSP cases (2) the false positive rates of the two sets of criteria in other FTLD cases (3) the clinical evolution of vPSP syndromes compared to PSP-RS and other syndromes associated with FTLD.

## METHODS

### Identification of cases

This work was approved by local institutional review boards and ethics committee at each participating center, according to their national legal and ethical requirements. Donors gave written informed consent before death according to the Declaration of Helsinki for the use of their brain tissue and medical records for research purposes; or next of kin provided consent after death, in the context of an *ante mortem* declaration of intention to donate, in accordance with each national law.

Patients with a neuropathological diagnosis of PSP^18, 19^ were identified from the neuropathology files of three brain banks with expertise in neurodegenerative diseases (Cambridge Brain Bank, Cambridge, UK; Neurological Tissue Bank of the Biobanc–Hospital Clinic-IDIBAPS, Barcelona, Catalonia, Spain; pathologically confirmed cases from the ENGENE-PSP study, USA, including cases from the University of Louisville, Maryland and Colorado). For comparison purposes, patients with a neuropathological diagnosis of CBD^19, 20^ and other FTLD (FTLD-tau, FTLD-TDP43, FTLD-DLDH), presenting either with behavioral variant frontotemporal dementia (bvFTD) or non-fluent/agrammatic variant of primary progressive aphasia (nfvPPA)^19, 21^, were collected from the Cambridge Brain Bank.

All cases were clinically evaluated by movement disorder specialists, behavioral neurologists, or psychiatrists with expertise in PSP, CBD and FTD, as documented in the patients’ files. Cases with insufficient clinical data were excluded. None of the patients in this study has been included in previous validation studies of PSP, although several cases of CBD were included in a prior study of CBD.^22^

### Data collection and clinical criteria

Retrospective chart review of the cases was performed to extract demographic and clinical information. ENGENE-PSP cases were prospectively evaluated as part of a case-control study, and data were collected at the first site Principal Investigator visit.^23^ In the whole sample, collected demographics included sex, estimated age at onset and age at first visit. For the Cambridge and Barcelona cohorts, age at death and disease duration were also collected. Standardized clinical features were recorded as previously described^17^, adding the feature of amantadine-induced hallucinations as distinct from levodopa induced hallucinations. These features were considered “present” if specifically mentioned in the clinical notes during the course of the disease, otherwise they were considered “absent”.

On the basis of the clinical phenotype, each PSP patient was classified according to the 1996 NINDS-SPSP and 2017 MDS-PSP criteria into the best fitting category (prioritizing early and predominant clinical features). Note that in the hierarchy of diagnostic certainty (probable>possible>suggestive), each case in the more certain category meets criteria for the lower category. Thus, probable-PSP cases all meet the lower criteria for possible-PSP. For cases from the Cambridge Brain Bank, the presence or absence of each feature was identified at the first visit, and when absent from the first visit, we recorded the timing of onset during the subsequent course of the disease. Thus, in this sub-cohort, it was possible to compare the two sets of criteria through the course of disease, and to use the retrospective clinical records to describe the clinical evolution of each syndrome as new symptoms and signs appeared. The same sets of criteria were applied to the other FTLD syndromes in order to assess the frequency of their clinical features.

### Statistical Analysis

Continuous variables were tested with either Mann-Whitney U test or Kruskal-Wallis test according to the number of groups compared. Categorical variables were tested with either Fisher’s exact test or Chi-square test according to the number of groups compared.

A p-value <0.05 was considered statistically significant for demographical variables. When testing clinical features, p values were adjusted following Bonferroni’s method to account for multiple comparisons (number of clinical features = 38; threshold p=0.0013).

For each set of criteria, sensitivity was defined as the percentage of cases fulfilling a clinical diagnosis of PSP among all PSP cases confirmed by neuropathological examination, while specificity as the percentage of cases without a clinical diagnosis of PSP among all non-PSP cases confirmed by neuropathological examination. False positive rate was defined as the percentage of cases fulfilling a clinical diagnosis of PSP among all the non-PSP cases confirmed by neuropathological examination.

## RESULTS

### 1.1. Demographic characteristics of the PSP cohort

108 neuropathologically confirmed cases of PSP were collected across the three sites (Cambridge: 65; Barcelona: 21; ENGENE-PSP: 22). Demographic data of the PSP cohort, according to site of origin, are reported in **Table 1**. There were no site differences with regard to sex, while age at onset and at first visit varied between sites. Post-hoc tests revealed that significance was driven by the comparison between samples from the ENGENE-PSP study and Barcelona (for age at onset) and comparison between Barcelona and both Cambridge and ENGENE-PSP study (for age at first visit).

**Table 1.** Demographic characteristics of the PSP and FTLD cohorts.

| **PSP cohort** |  |  |  |  |
| --- | --- | --- | --- | --- |
|  | *Cambridge (N=65)* | *Barcelona (N=21)* | *ENGINE-PSP (N=22)* | *p* |
| Sex, female, % | 43.1 | 61.9 | 45.5 | n.s.<sup>a</sup> |
| Age at onset, yrs | 67.0±7.2 | 69.8±6.5 | 63.6±6.2 | 0.024* |
| Age at first visit, yrs | 69.8±7.1 | 74.7±4.5 | 67.3±6.4 | 0.001<sup>*,†</sup> |
| **FTLD cohort** |  |  |  |  |
|  | *bvFTD (N=38)* | *nfvPPA (N=14)* | *CBD (N=29)* | *p* |
| Sex, female, % | 23.7 | 28.6 | 41.4 | n.s.<sup>a</sup> |
| Age at onset, yrs | 58.3±9.1 | 61.1±8.6 | 64.3±6.9 | 0.017<sup>^</sup> |
| Age at first visit, yrs | 61.7±9.3 | 63.6±7.8 | 66.9±6.8 | 0.04<sup>^</sup> |
| Age at death, yrs | 67.5±10.1 | 70.1±8.3 | 71.6±7.1 | n.s. |
| Disease duration, yrs | 9.2±4.6 | 9.0±3.5 | 7.3±3.9 | n.s. |
| FTLD-Tau | 42.1 (16) | 50.0 (7) |  |  |
| FTLD-TDP-43 | 44.7 (17) | 42.9 (6) |  |  |
| FTLD-DLDH | 13.2 (5) | 7.1 (1) |  |  |
| FTLD-FUS | 0 (0) | 0 (0) |  |  |

Kruskal-Wallis test, otherwise specified: <sup>a</sup>Chi-square test. Results are expressed as mean ± standard deviation. Sex is expressed as percentage.
\* refers to Barcelona vs ENGINE-PSP comparison (p<0.05 post-hoc comparison).
<sup>†</sup> refers to Barcelona vs Cambridge comparison (p<0.05 post-hoc comparison).
<sup>^</sup> refers to bvFTD vs CBD comparison (p<0.05 post-hoc comparison).
yrs: years; n.s.: not significant; FTLD: frontotemporal lobar degeneration; bvFTD: behavioural variant of frontotemporal dementia; nfvPPA: non-fluent/agrammatic variant of primary progressive aphasia; CBD: corticobasal degeneration; TDP-43: TAR DNA-binding protein 43; DLDH: dementia lacking distinctive histologic features; FUS: fused in sarcoma protein.

### 1.2. Sensitivity, specificity and false positive rates of the new MDS-PSP criteria and NINDS-SPSP criteria

The 2017 MDS-PSP criteria demonstrated high sensitivity for identification of PSP pathology (72.2% if “probable”, 100% if “suggestive of”), compared to medium sensitivity of NINDS-SPSP criteria (48.1-61.1%), when applied at the first assessment. Conversely, specificity was higher for NINDS-SPSP criteria (97.5-100%) than for MDS-PSP criteria (53.1-95.1%). Notably, the low specificity of MDS-PSP criteria was mainly due to false positive diagnosis at the “suggestive of” level of certainty, being otherwise good for “possible” and “probable” definitions (88.9-95.1%) (**Table 2**).

**Table 2.**
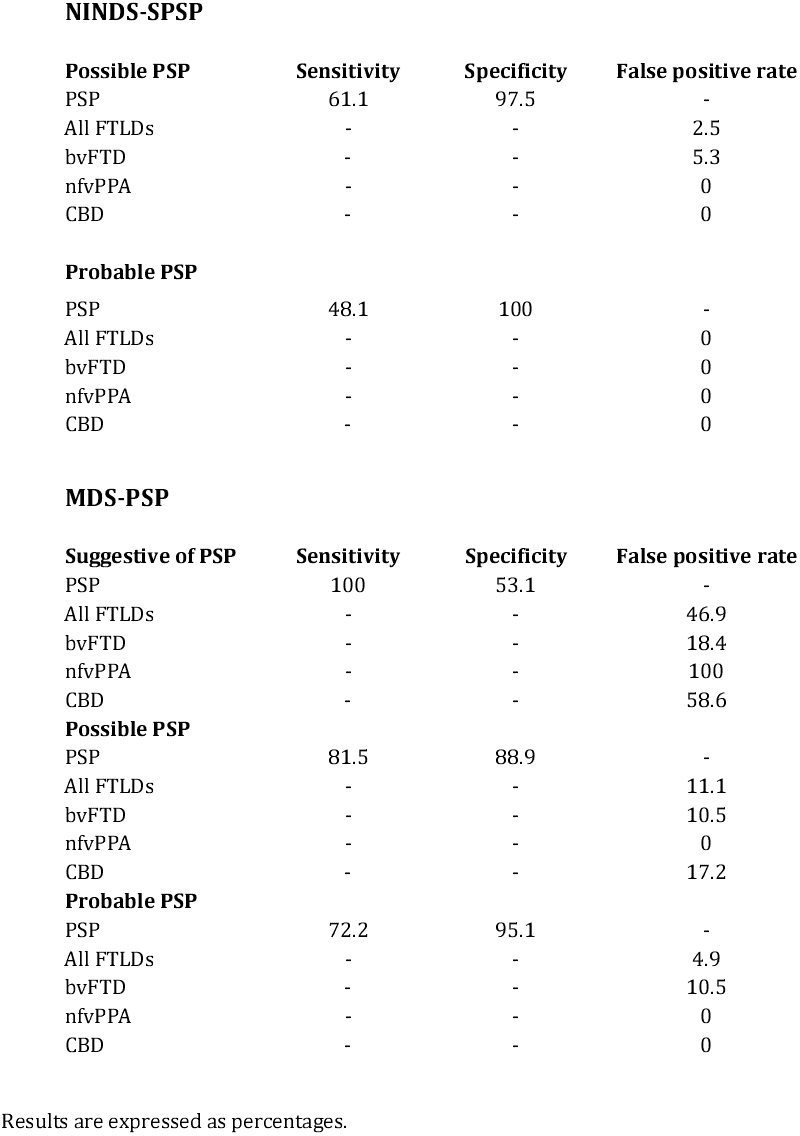
Sensitivity, specificity and false positive rates for each set of criteria and each degree of certainty.

**NINDS-SPSP**
| **Possible PSP** | **Sensitivity** | **Specificity** | **False positive rate** |
| --- | --- | --- | --- |
| PSP | 61.1 | 97.5 | - |
| All FTLDs | - | - | 2.5 |
| bvFTD | - | - | 5.3 |
| nfvPPA | - | - | 0 |
| CBD | - | - | 0 |
| **Probable PSP** |  |  |  |
| PSP | 48.1 | 100 | - |
| All FTLDs | - | - | 0 |
| bvFTD | - | - | 0 |
| nfvPPA | - | - | 0 |
| CBD | - | - | 0 |

**MDS-PSP**
| **Suggestive of PSP** | **Sensitivity** | **Specificity** | **False positive rate** |
| --- | --- | --- | --- |
| PSP | 100 | 53.1 | - |
| All FTLDs | - | - | 46.9 |
| bvFTD | - | - | 18.4 |
| nfvPPA | - | - | 100 |
| CBD | - | - | 58.6 |
| **Possible PSP** |  |  |  |
| PSP | 81.5 | 88.9 | - |
| All FTLDs | - | - | 11.1 |
| bvFTD | - | - | 10.5 |
| nfvPPA | - | - | 0 |
| CBD | - | - | 17.2 |
| **Probable PSP** |  |  |  |
| PSP | 72.2 | 95.1 | - |
| All FTLDs | - | - | 4.9 |
| bvFTD | - | - | 10.5 |
| nfvPPA | - | - | 0 |
| CBD | - | - | 0 |

Results are expressed as percentages.

Indeed, 61.1% (N=66) of patients fulfilled NINDS-SPSP criteria (48.1% being “probable” PSP), while 38.9% (N=42) did not satisfy either definition (**Table 2**). With the new MDS-PSP criteria, 59.3% (N=64) of patients fulfilled criteria for PSP-RS, followed by PSP-P (10.2%, N=11), PSP-CBS (9.3%, N=10), PSP-SL (9.3%, N=10), PSP-F (6.5%, N=7), PSP-PI (3.7%, N=4), PSP-PGF (0.9%, N=1) and PSP-OM (0.9%, N=1). None of the patients was misclassified (see **Supplementary Table 1**, note that in this table all “probable” cases meet the lower criteria for “possible” and “suggestive of” PSP). Of these subjects, 72.2% (N=78) of patients fell into the most stringent category “probable”, while 81.5% (N=88) met criteria for “possible” and 100% (N=108) met criteria for the “suggestive of” category (**Table 2**).

Just 2 out of 81 of the other FTLD cases fulfilled NINDS-PSP criteria for “possible” PSP, leading to a false positive rate of 2.5%. Looking at each of the FTLD conditions, false positive rate with NINDS-SPSP criteria was 5.3 for bvFTD (N=2) and nil for CBD and nfvPPA. None of them fulfilled “probable” criteria. Note that when these cases were clinically diagnosed as having other disorders than PSP, the critical clinical features of falls and vertical gaze palsy were late or minor in relation to the other clinical features (whereas falls occur <1 year under NINDS-SPSP and <3 years under MDS-PSP).

In contrast, 38 of the other 81 FTLD cases fulfilled one of the MDS-PSP definitions, leading to a false positive rate of 46.9%. When looking at each degree of certainty, false positive rate ranged from 46.9% (38 out of 81) for “suggestive of”, to 11.1% (9 out of 81) for “possible” and 4.9% (4 out of 81) for “probable”. While the “suggestive of” definitions did not allow to correctly identify the underlying pathology in nfvPPA and CBD cases (with a false positive rate of 100 and 58.6%, respectively), it was low for bvFTD cases (18.4%) (**Table 2** and **Supplementary Table 2**). More details about sensitivity and specificity in each cohort are reported in **Supplementary Table 3**.

### 2.1. Demographic characteristics of PSP-RS and vPSP cases

Of the 65 PSP cases from the Cambridge cohort, at the first visit 50.8% (N=33) fulfilled clinical criteria for probable PSP-RS, while 49.2% (N=32) for vPSP. Among vPSP cases, PSP-SL was the most frequent presentation (15.4%, N=10), followed by PSP-CBS (12.3%, N=8), PSP-F (7.7%, N=5), PSP-P (6.1%, N=4), PSP-PI (6.1%, N=4) and PSP-OM (1.5%, N=1). vPSP cases were older at death compared to PSP-RS cases (p=0.041), but with a comparable disease duration (noting that this group is heterogeneous and we lack power to separate the vPSP syndromes). There were no differences with regard to sex distribution. Demographic variables of these cohorts are reported in **Table 3**.

**Table 3.** Demographic characteristics of PSP-RS and vPSP cases.

|  | **PSP-RS (N=33)** | **vPSP (N=32)** | **p** |
| --- | --- | --- | --- |
| Sex, female, % | 45.5 | 40.6 | n.s.<sup>a</sup> |
| Age at onset, yrs | 65.6±6.7 | 68.4±7.6 | n.s. |
| Age at first visit, yrs | 68.6±6.5 | 71.2±7.6 | n.s. |
| Age at death, yrs | 72.3±6.3 | 75.4±7.3 | 0.041 |
| Disease duration, yrs | 6.7±2.4 | 7.0±3.1 | n.s. |

Mann-Whitney U test, otherwise specified: <sup>a</sup>Fisher's exact test. Results are expressed as mean ± standard deviation. Sex is expressed as percentage.
PSP-RS: PSP with Richardson's syndrome; vPSP: variant PSP syndromes; n.s.: not significant.

### 2.2. Evolution of PSP features in PSP-RS and vPSP cases

In keeping with the application of the diagnostic criteria, at first visit PSP-RS presented with higher prevalence of falls (p<0.001), abnormal saccades or pursuits (p<0.001), supranuclear gaze palsy (p<0.001) and prevalent axial rigidity (p=0.002), while vPSP presented more frequently with non-fluent aphasia (p<0.001). However, by the last visit, the prevalence of these cardinal PSP-RS features was similar. The rising prevalence of each feature over the years from diagnosis is illustrated in **Figure 1** and compared between first and last visit in **Supplementary Table 4**.

**Figure 1.**
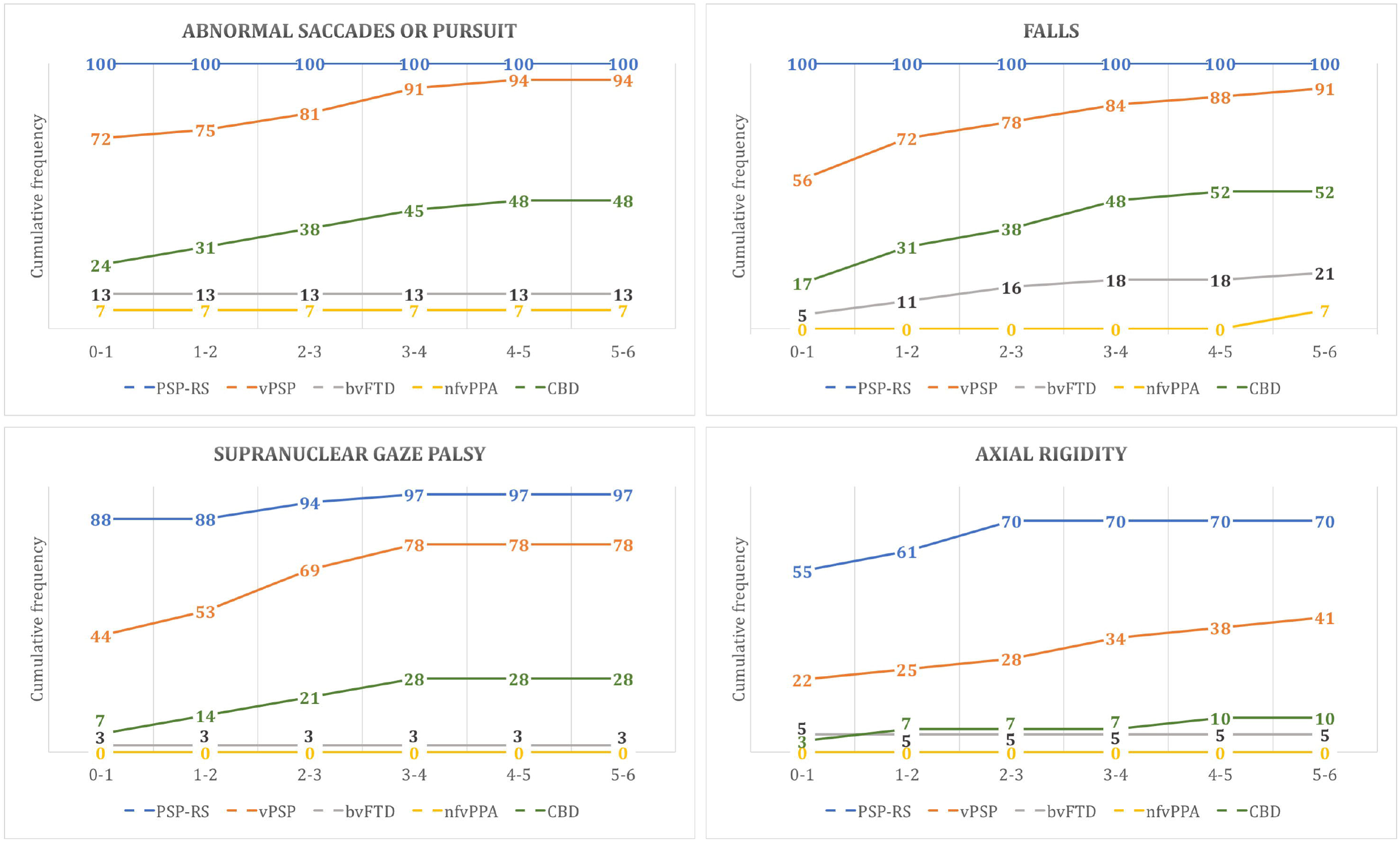
Cumulative frequency of typical PSP-RS signs/symptoms. Line charts reporting the cumulative frequency of significantly overexpressed signs /symptoms at first visit in the PSP-RS cohort compared to vPSP and each non-PSP neuropathological cohort. Time is reported on the x-axis at yearly intervals from the first visit. Cumulative frequency is reported as percentage on the y-axis.

### 3.1. Demographic characteristics of PSP-F, PSP-CBS, PSP-SL and the corresponding FTLD syndromes

Among 32 vPSP cases, 5 fulfilled MDS clinical criteria for PSP-F, 8 for PSP-CBS and 10 for PSP-SL. From the same brain bank, 81 cases fulfilled neuropathological criteria for FTLD other than PSP, with either frontotemporal dementia (38 bvFTD, 14 nfvPPA phenotypes) or CBD (N=29). Of the 38 bvFTD cases, 44.7% (N=17) presented with TDP-43 pathology, 42.1% (N=16) with tau pathology and 13.2% (N=5) with dementia lacking distinctive histologic features. Of the 14 nfvPPA cases, 50.0% (N=7) presented with tau pathology, 42.9% (N=6) with TDP-43 pathology and 7.1% (N=1) with dementia lacking distinctive histologic features (**Table 1**).

PSP-F presented shorter disease duration (mean 4.6±0.7) compared to bvFTD cases, despite comparable age at onset and age at death. In contrast, PSP-CBS and PSP-SL were significantly older than CBD and nfvPPA cases, but with an overall comparable disease duration (see **Table 4**). There were no differences with regard to sex distribution.

**Table 4.** Demographic characteristics of PSP-F, PSP-CBS, PSP-SL and the corresponding FTLD syndromes.

|  | PSP-F (N=5) | bvFTD (N=38) | p | PSP-CBS (N=8) | CBD (N=29) | p | PSP-SL (N=10) | nvPPA (N=14) | p |
| --- | --- | --- | --- | --- | --- | --- | --- | --- | --- |
| Sex, female, % | 0 (0) | 23.7 (9) | n.s.<sup>a</sup> | 50.0 (4) | 41.4 (12) | n.s.<sup>a</sup> | 60.0 (6) | 28.6 (4) | n.s.<sup>a</sup> |
| Age at onset, yrs | 63.7±7.0 | 58.3±9.1 | n.s. | 73.9±5.7 | 64.3±6.9 | 0.002 | 69.4±3.6 | 61.1±8.6 | 0.006 |
| Age at first visit, yrs | 65.9±7.1 | 61.7±9.3 | n.s. | 77.6±6.6 | 66.9±6.8 | 0.001 | 71.5±3.2 | 63.6±7.8 | 0.005 |
| Age at death, yrs | 68.3±7.1 | 67.5±10.1 | n.s. | 80.4±5.7 | 71.6±7.1 | 0.002 | 76.9±3.2 | 70.1±8.3 | 0.016 |
| Disease duration, yrs | 4.6±0.7 | 9.2±4.6 | 0.006 | 6.5±3.8 | 7.3±3.9 | n.s. | 7.5±2.4 | 9.0±3.5 | n.s. |

Mann-Whitney U test, otherwise specified: <sup>a</sup>Fisher's exact test. Results are expressed as mean ± standard deviation. Sex is expressed as percentage.
PSP-F: PSP with predominant frontal presentation; PSP-CBS: PSP with predominant corticobasal syndrome; PSP-SL: PSP with predominant speech/language disorder.

### 3.2. Evolution of features in vPSP and their corresponding FTD/CBD syndromes

#### PSP-F vs bvFTD

At the first visit, PSP-F presented higher prevalence of abnormal saccades or pursuit (p=0.003), dysarthria (p=0.024) and supranuclear gaze palsy (p=0.032), than those with bvFTD presentations of FTLD other than PSP. However, none of the reported features survived correction for multiple comparisons. At last visit, abnormal saccades/pursuit (p<0.001), bradykinesia (p<0.001), supranuclear gaze palsy (p<0.001), postural instability (p=0.001) and dysphagia (p=0.001) were significantly overexpressed in PSP-F cases. None of the tested features was significantly more present in bvFTD cases (see **Supplementary Table 5** for an extended report).

#### PSP-CBS vs CBD

At the first visit, falls were significantly more recorded in PSP-CBS cases than in patients with CBD associated with non-PSP phenotypes (p=0.001). Limb dystonia (p=0.005), non-specific visual symptoms (p=0.005) and bradykinesia (p=0.008) were more common in PSP-CBS cases, but these differences did not survive the threshold for Bonferroni corrected statistics. At the last visit, none of the recorded features was significantly over-expressed in either group (see **Supplementary Table 6** for an extended report).

#### PSP-SL vs nfvPPA

No specific feature discriminated between PSP-SL and nfvPPA cases at the first visit, while at the last visit the presence of falls, abnormal saccades or pursuit and supranuclear gaze palsy were significantly more common in the PSP-SL population (p<0.001) (see **Supplementary Table 7** for an extended report).

## DISCUSSION

This independent cohort of neuropathologically-proven cases of PSP, CBD and other FTLD phenotypes, shows that at the first specialist assessment the revised MDS-PSP clinical diagnostic criteria succeeded in their aims of high sensitivity both to the classical PSP-RS presentation and to multiple variant phenotypes of PSP. In addition, longitudinal assessment reveals the convergence from vPSP phenotypes towards PSP-RS, with emergence of supranuclear gaze palsy and falls. For disease-modifying therapies for PSP and related disorders, the correct identification of candidate subjects in the early stages of the disease has become pressing. In the absence of sufficiently sensitive and specific non-invasive biomarkers to discriminate PSP from other parkinsonian and behavioral disorders^24–28^, diagnosis still relies on clinical evaluation. The ability of clinical diagnostic criteria to identify and differentiate PSP is therefore critical.^15^ We suggest that the new criteria are sufficiently sensitive at relatively early stages of disease (at least, when first assessed by specialist teams) to consider using for selection of cases for novel PSP-therapies, and among vPSP the probable and possible criteria are sufficiently specific to differentiate PSP-F from bvFTD.

The MDS-PSP criteria for “suggestive of” (including also patients meeting more stringent criteria for possible/probable PSP) reached 100% sensitivity, compared to 61.1% of NINDS-SPSP “possible” criteria. This improvement was driven partly by less restrictive criteria for PSP-RS (with falls being considered significant within three years instead of one), inclusion of suggestive criteria and partly by the recognition of the variant syndromes. As a consequence of more liberal and diverse criteria by MDS-PSP, there was a risk that neuropathologically-confirmed cases of FTLD (FTLD-tau, FLD-TDP43, CBD) with syndromes of bvFTD, nfvPPA and CBS, could be misdiagnosed as PSP. Despite this possibility, the MDS-PSP criteria succeed in offering a more comprehensive clinical spectrum of PSP, with operationalization of the diagnoses where there is clinical suspicion of possible underlying PSP. In addition to “probable” and “possible” categories, the new criteria introduce the category of “suggestive”. This category has been designed to recognize early signs of PSP which do not meet threshold for possible or probable PSP. Such patients require close clinical follow-up to identify progression to either PSP or to a non-PSP phenotype^15, 16^, and call for the development of better biomarkers to ascertain tau pathology at early stages of symptomatic disease.^16^ In line with this, specificity of MDS-PSP criteria increased significantly when not including cases that were only “suggestive” (and which did not meet more stringent criteria for possible/probable), which were highly affected by false recognition of nfvPPA and CBD cases. This study also characterized the clinical presentation and evolution of vPSP cases compared to PSP-RS. Specialist clinics vary in the proportion of PSP-RS and vPSP cases they evaluate, reflecting in part their emphasis on movement *versus* cognitive disorders. Whilst the exact proportion varies, it is now clear that vPSP cases constitute a substantial proportion of the burden of PSP pathology.^11^ In our cohort, PSP-SL was the most frequent variant presentation, followed by PSP-CBS and PSP-F. In a previous study of vPSP, PSP-SL was less frequently described, and PSP-P was more common.^11^ These differences in distribution of vPSP types may arise from differential referral patterns in neurological services, for example to cognitive disorder clinics, movement disorder clinics, and integrated clinical services.

Overall, vPSP cases showed similar age at onset and disease duration compared to PSP-RS, and the only discriminative features at onset were represented by typical PSP-RS symptoms (i.e. falls and oculomotor disturbances), which develop later in the course for all patients with vPSP presentations. This indicates a convergence of phenotypes.^8, 11, 29^

Despite the relatively small number of vPSP cases, we examined the clinical evolution of the vPSP phenotypes (i.e. PSP-SL, PSP-CBS and PSP-F). PSP-F cases showed a more aggressive disease course than bvFTD cases, with a mean survival of 4.6 years from onset. PSP-CBS and PSP-SL presented similar disease duration to CBD (of any phenotype) and nfvPPA (with FTLD pathology) respectively, despite being older at onset. Dementia, dysphagia and supranuclear gaze palsy have been variably reported to be clinical predictors of death in PSP-RS.^30, 31^ In our PSP-F cohort, both dysphagia and supranuclear gaze palsy were over-expressed compared to bvFTD patients, suggesting a possible role of either symptom on survival. However, dysphagia was the only symptom not consistently over-expressed in PSP-SL and PSP-CBS cases. Although not a primary objective of this study, these results raise the hypothesis that early dysphagia account for higher mortality in PSP-F patients.^31, 32^

At onset, the prevalence of many recorded features was similar between groups. PSP-RS-indicators became prominent later in the course of vPSP syndromes, and were significantly more common only in PSP-F and PSP-SL cases. The PSP-CBS and CBD cases showed clinical overlap across all the disease course, underlining the well-known neuropathological similarities between PSP and CBD and borderline cases.^33, 34^ Nonetheless, the onset of RS-like features even in atypical cases raise the likelihood of underlying 4-repeat tau pathology.^29, 35^ Despite these encouraging findings, we acknowledge some limitations. First of all, because of the retrospective nature of the study, clinical assessment and report of features was not standardized across sites, although there was within-site standardization (e.g. EUGENE-PSP proformas). Consequently, the documented clinical data may have varied with regard to its level of detail and focus. In keeping with the development study dataset for the MDS-PSP criteria, some cognitive and behavioral features were collapsed together into super-ordinate categories (i.e. “frontal lobe dysfunction”, “executive dysfunction”, “frontal personality changes”, “frontal social dysfunction”) in order to reduce the number of variables tested, but this obviously comes with the limitation of allowing less precision with regard to the PSP cognitive/behavioral profile and differences with other conditions. Furthermore, this study lacks comparison with pathologically proven cases of Parkinson’s disease and multiple system atrophy. However, one of the major structural changes between NINDS-SPSP and MDS-PSP was the addition of the criteria for PSP variants with language, behavioral and corticobasal presentations. These raised the problem of specificity with respect to the other FTLD pathologies presenting with behavioural (bvFTD), language (nfvPPA) and corticobasal (CBS) syndromes. The current study was oriented to this particular challenge. A second structural change in the MDS-PSP criteria was the introduction of the category “suggestive of” PSP syndromes. Whilst we included such cases in our analysis, we recognize that many people with problems “suggestive of” PSP may present through other healthcare services (e.g. ophthalmology, falls clinics) without reaching specialist movement disorder or cognitive neurology services. Identification and inclusion of such people in long-term research studies and pathology is required.

In conclusion, our study provides good validation of the 2017 MDS-PSP criteria and delineates the clinical evolution of vPSP cases with neuropathological confirmation. The new criteria are promising for identification of both PSP-RS and vPSP, with the expected occasional misdiagnosis with other FTLD pathologies. Using the lowest certainty criteria (“suggestive of”), bvFTD was falsely recognized by the MDS-PSP criteria at first assessment at a low rate (18.4%), while 58% of CBD and 100% of nfvPPA were “suggestive of’ PSP. The application of stricter “possible” and “probable” criteria led to the complete discrimination of PPA cases due to non-PSP type FTLD pathologies. In terms of potential treatment based on likely PSP pathology, our results suggest that particular care must be taken regarding the inclusion of PPA cases in clinical trials, as the MDS-PSP criteria did not discriminate between the neuropathological substrates of the PSP-SL phenotype. The inclusion of misidentified CBD cases may not pose such a problem for treatments targeting the common features of PSP and CBD pathologies (i.e. both 4-repeat tau disorders). In this context, the development of biomarkers with high specificity (such as tau-specific imaging) may further improve accuracy in early stages. The application of the new MDS-PSP criteria will further facilitate the development of such tools. Prospective studies with systematic data collection will be useful to corroborate these findings, but we tentatively suggest that clinical triallists can proceed with confidence in the application of the MDS-PSP criteria.

## AKNOWLEDGMENTS

To all the patients and their next of kin for their goodwill and generosity, as without them none of this brain-bank based research would have been possible. We are also grateful to all the neurologists and other physicians and health professionals who carefully and proficiently assessed the clinical features and diagnoses of the brain donors and eventually obtained their consent for brain donation.

## AUTHORS’ ROLES

Stefano Gazzina - acquisition and interpretation of data, statistical analysis, drafting of original manuscript.

Gesine Respondek - critical revision of manuscript for important intellectual content.

Yaroslau Compta - acquisition and interpretation of data, critical revision of manuscript for important intellectual content.

Kieren S.J. Allinson – critical revision of manuscript for important intellectual content.

Maria G. Spillantini – critical revision of manuscript for important intellectual content.

Laura Molina-Porcel – acquisition and interpretation of data, critical revision of manuscript for important intellectual content.

Mar Guasp-Verdaguer – acquisition and interpretation of data, critical revision of manuscript for important intellectual content.

Shirin Moftakhar - acquisition and interpretation of data.

Stephen G. Reich - acquisition and interpretation of data, critical revision of manuscript for important intellectual content.

Deborah Hall - acquisition and interpretation of data, critical revision of manuscript for important intellectual content.

Irene Litvan - acquisition and interpretation of data, critical revision of manuscript for important intellectual content.

Günter U. Höglinger - critical revision of manuscript for important intellectual content.

James B. Rowe - study concept and design, study supervision, drafting of original manuscript, critical revision of manuscript for important intellectual content.

## FINANCIAL DISCLOSURES

Stefano Gazzina – None

Gesine Respondek – has received honoraria from GE Healthcare.

Yaroslau Compta – has received funding, research support and/or honoraria from and has served on advisory boards for UCB pharma, Lundbeck, Medtronic, Abbvie, Novartis, GSK, Boehringer, Pfizer, Merz, Teva, Zambon, Bial, Ipsen, Piramal Imaging and Esteve. He has also received research funding from the H2020 EU programme, the European Regional Development Fund (through the ISCIII), and the CERCA Programme of the Generalitat de Catalunya.

Kieren S.J. Allinson –

Maria G. Spillantini –

Laura Molina-Porcel – None

Mar Guasp-Verdaguer – None

Shirin Moftakhar – None

Stephen G. Reich –

Deborah Hall –

Irene Litvan – was a member of the Abbie Advisory Board, consultant for Toyama Pharmaceuticals and member of the Biotie/Parkinson Study Group Medical and Biogen steering scientific committees. Her research is supported by the National Institutes of Health grants: 5P50 AG005131-31, 5T35HL007491,1U01NS086659, 1U54NS092089-01, and U01NS100610; Parkinson Study Group, Michael J Fox Foundation, AVID Pharmaceuticals, C2N Diagnostics and Bristol-Myers Squibb. She receives her salary from the University of California San Diego and as Editor of Frontiers in Neurology.

Günter U. Höglinger – served on the advisory boards for AbbVie, Alzprotect, Asceneuron, Bristol-Myers Squibb, Biogen, Novartis, Roche, Sellas Life Sciences Group, UCB; has received honoraria for scientific presentations from Abbvie, Biogen, Roche, Teva, UCB, has received research support from CurePSP, the German Academic Exchange Service (DAAD), German Parkinson’s Disease Foundation (DPG), German PSP Association (PSP Gesellschaft), German Research Foundation (DFG) and the German Ministry of Education and Research (BMBF), International Parkinson’s Fonds (IPF), Neuropore, the Sellas Life Sciences Group; has received institutional support from the German Center for Neurodegenerative Diseases (DZNE).

James B. Rowe – is supported by the Wellcome Trust (103838) and the NIHR Cambridge Biomedical Research Centre. Additional grant funding is received from the Medical Research Council, the McDonnell Foundation, Alzheimer’s Research UK, Evelyn Trust, Dementias Platform UK, and research grants from AZ-Medimmune and Eli Lilly. He is a consultant for Asceneuron and serves as Editor for the journal Brain.

## SUPPLEMENTARY MATERIAL

**Supplementary Table 1.** Sensitivity of the two sets of criteria in the whole PSP cohort.

| **NINDS-SPSP** |  |  |
| --- | --- | --- |
|  | **N** | **Percentage** |
| Probable PSP# | 52 | 48.1 |
| Possible PSP only | 14 | 13.0 |
| Unclassified | 42 | 38.9 |
| Sensitivity |  | 61.1 |
| **MDS-PSP** |  |  |
|  | **N** | **Percentage** |
| Probable PSP-RS# | 64 | 59.3 |
| Suggestive of PSP-SL | 8 | 7.4 |
| Probable PSP-P# | 7 | 6.5 |
| Probable PSP-F# | 6 | 5.6 |
| Possible PSP-CBS# | 6 | 5.6 |
| Suggestive of PSP-P | 4 | 3.7 |
| Suggestive of PSP-CBS | 4 | 3.7 |
| Suggestive of PSP-PI | 4 | 3.7 |
| Possible PSP-SL# | 2 | 1.9 |
| Possible PSP-F# | 1 | 0.9 |
| Probable PSP-PGF# | 1 | 0.9 |
| Possible PSP-OM# | 1 | 0.9 |
| Unclassified | 0 | 0 |
| Sensitivity |  | 100 |

For each diagnostic category, the number and the percentage of subjects fulfilling the definition are reported.

**Supplementary Table 2.**
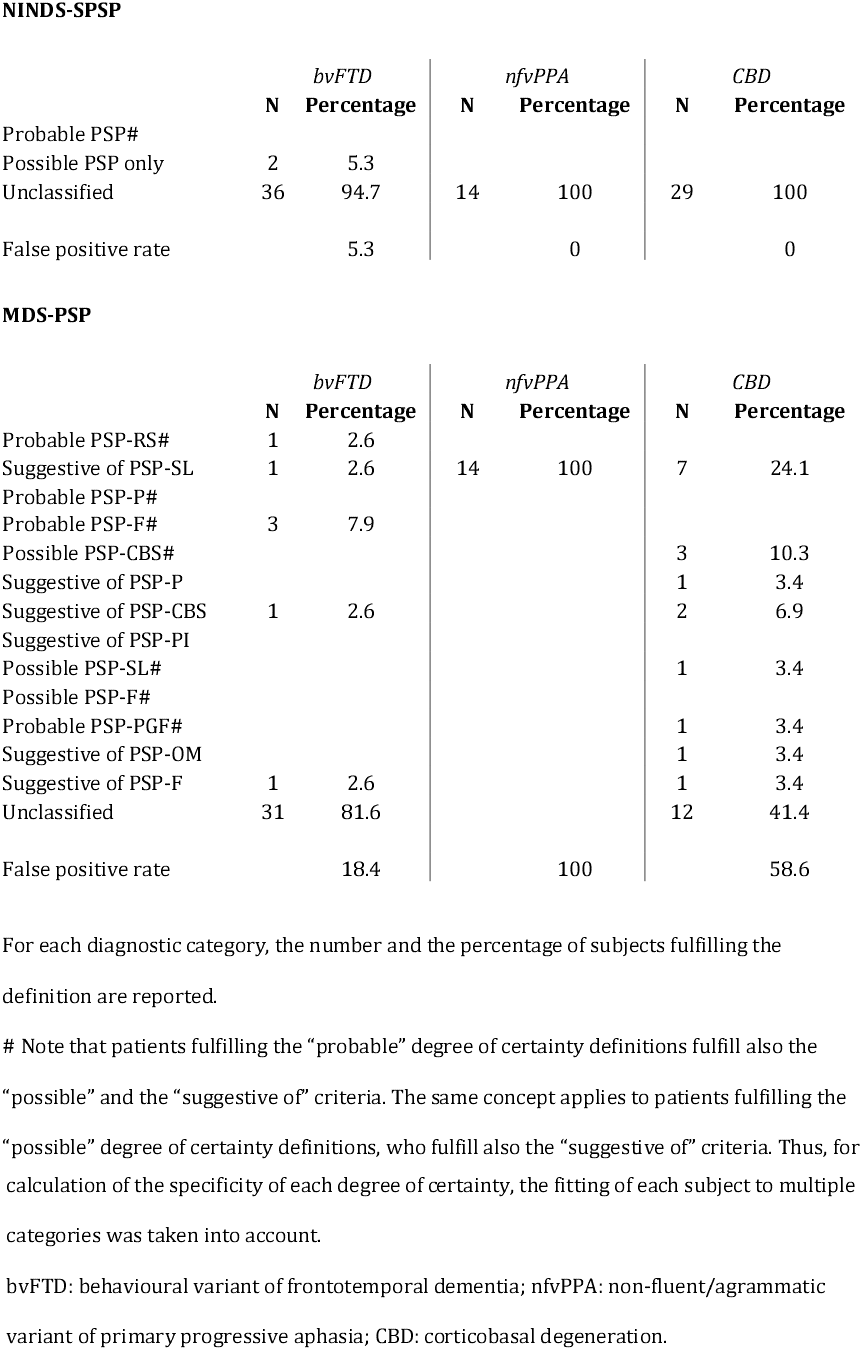
False positive rate of the two sets of criteria in the separate FTLD cohorts.

**NINDS-SPSP**
|  | *bvFTD* |  | *nfvPPA* |  | *CBD* |  |
| --- | --- | --- | --- | --- | --- | --- |
|  | N | Percentage | N | Percentage | N | Percentage |
| Probable PSP# |  |  |  |  |  |  |
| Possible PSP only | 2 | 5.3 |  |  |  |  |
| Unclassified | 36 | 94.7 | 14 | 100 | 29 | 100 |
| False positive rate |  | 5.3 |  | 0 |  | 0 |

**MDS-PSP**
|  | *bvFTD* |  | *nfvPPA* |  | *CBD* |  |
| --- | --- | --- | --- | --- | --- | --- |
|  | N | Percentage | N | Percentage | N | Percentage |
| Probable PSP-RS# | 1 | 2.6 |  |  |  |  |
| Suggestive of PSP-SL | 1 | 2.6 | 14 | 100 | 7 | 24.1 |
| Probable PSP-P# |  |  |  |  |  |  |
| Probable PSP-F# | 3 | 7.9 |  |  |  |  |
| Possible PSP-CBS# |  |  |  |  | 3 | 10.3 |
| Suggestive of PSP-P |  |  |  |  | 1 | 3.4 |
| Suggestive of PSP-CBS | 1 | 2.6 |  |  | 2 | 6.9 |
| Suggestive of PSP-PI |  |  |  |  |  |  |
| Possible PSP-SL# |  |  |  |  | 1 | 3.4 |
| Possible PSP-F# |  |  |  |  |  |  |
| Probable PSP-PGF# |  |  |  |  | 1 | 3.4 |
| Suggestive of PSP-OM |  |  |  |  | 1 | 3.4 |
| Suggestive of PSP-F | 1 | 2.6 |  |  | 1 | 3.4 |
| Unclassified | 31 | 81.6 |  |  | 12 | 41.4 |
| False positive rate |  | 18.4 |  | 100 |  | 58.6 |

For each diagnostic category, the number and the percentage of subjects fulfilling the definition are reported.

**Supplementary Table 3.** Sensitivity of the two sets of criteria in the separate PSP cohorts.

**NINDS-SPSP**
|  | *Cambridge* |  | *Barcelona* |  | *ENGINE-PSP* |  |
| --- | --- | --- | --- | --- | --- | --- |
|  | **N** | **Percentage** | **N** | **Percentage** | **N** | **Percentage** |
| Probable PSP# | 29 | 44.6 | 2 | 9.5 | 21 | 95.5 |
| Possible PSP only | 9 | 13.9 | 4 | 19.1 | 1 | 4.5 |
| Unclassified | 27 | 41.5 | 15 | 71.4 |  |  |
| Sensitivity |  | 58.5 |  | 28.6 |  | 100 |

**MDS-PSP**
|  | *Cambridge* |  | *Barcelona* |  | *ENGINE-PSP* |  |
| --- | --- | --- | --- | --- | --- | --- |
|  | **N** | **Percentage** | **N** | **Percentage** | **N** | **Percentage** |
| Probable PSP-RS# | 33 | 50.8 | 9 | 42.9 | 22 | 100 |
| Suggestive of PSP-SL | 8 | 12.3 |  |  |  |  |
| Probable PSP-P# | 1 | 1.5 | 6 | 28.6 |  |  |
| Probable PSP-F# | 5 | 7.7 | 1 | 4.8 |  |  |
| Possible PSP-CBS# | 5 | 7.7 | 1 | 4.8 |  |  |
| Suggestive of PSP-P | 3 | 4.6 | 1 | 4.8 |  |  |
| Suggestive of PSP-CBS | 3 | 4.6 | 1 | 4.8 |  |  |
| Suggestive of PSP-PI | 4 | 6.2 |  |  |  |  |
| Possible PSP-SL# | 2 | 3.1 |  |  |  |  |
| Possible PSP-F# |  |  | 1 | 4.8 |  |  |
| Probable PSP-PGF# |  |  | 1 | 4.8 |  |  |
| Possible PSP-OM# | 1 | 1.5 |  |  |  |  |
| Sensitivity |  | 100 |  | 100 |  | 100 |

For each diagnostic category, the number and the percentage of subjects fulfilling the definition are reported.

**Supplementary Table 4.**
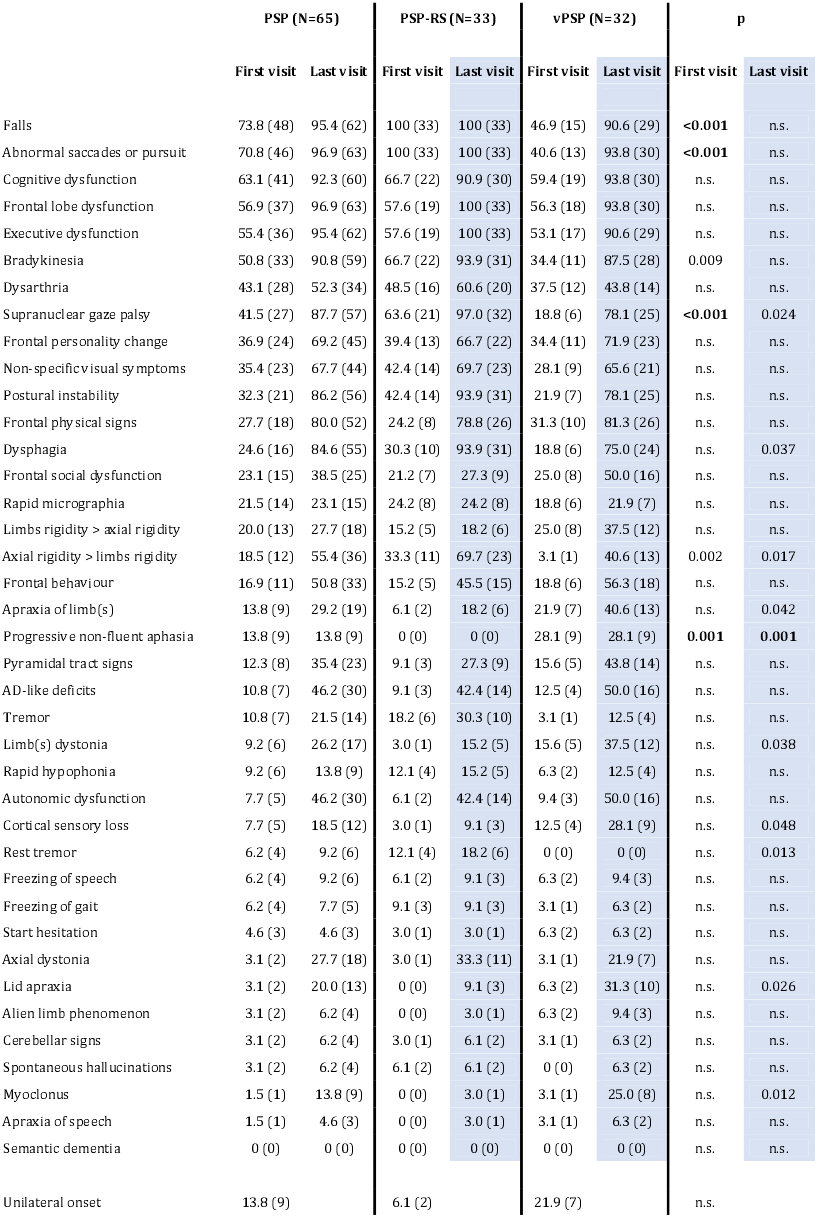
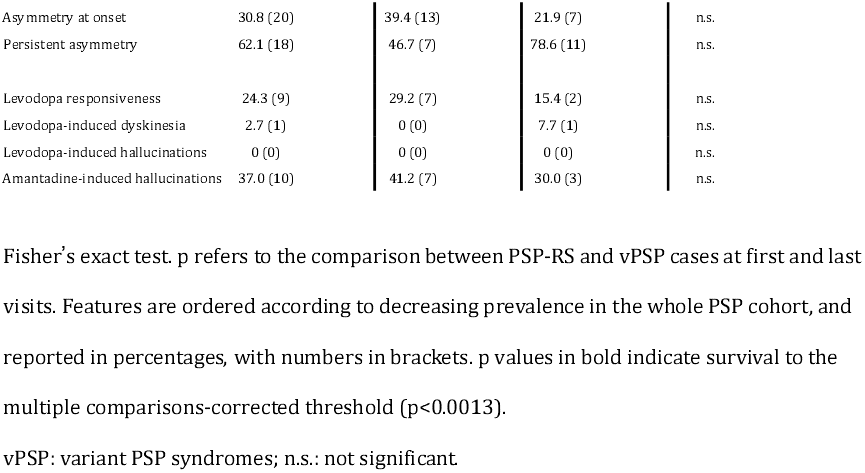
Prevalence of PSP features in PSP-RS and vPSP cases.

|  | PSP (N=65) |  | PSP-RS (N=33) |  | vPSP (N=32) |  | p |  |
| --- | --- | --- | --- | --- | --- | --- | --- | --- |
|  | First visit | Last visit | First visit | Last visit | First visit | Last visit | First visit | Last visit |
| Falls | 73.8 (48) | 95.4 (62) | 100 (33) | 100 (33) | 46.9 (15) | 90.6 (29) | **<0.001** | n.s. |
| Abnormal saccades or pursuit | 70.8 (46) | 96.9 (63) | 100 (33) | 100 (33) | 40.6 (13) | 93.8 (30) | **<0.001** | n.s. |
| Cognitive dysfunction | 63.1 (41) | 92.3 (60) | 66.7 (22) | 90.9 (30) | 59.4 (19) | 93.8 (30) | n.s. | n.s. |
| Frontal lobe dysfunction | 56.9 (37) | 96.9 (63) | 57.6 (19) | 100 (33) | 56.3 (18) | 93.8 (30) | n.s. | n.s. |
| Executive dysfunction | 55.4 (36) | 95.4 (62) | 57.6 (19) | 100 (33) | 53.1 (17) | 90.6 (29) | n.s. | n.s. |
| Bradykinesia | 50.8 (33) | 90.8 (59) | 66.7 (22) | 93.9 (31) | 34.4 (11) | 87.5 (28) | 0.009 | n.s. |
| Dysarthria | 43.1 (28) | 52.3 (34) | 48.5 (16) | 60.6 (20) | 37.5 (12) | 43.8 (14) | n.s. | n.s. |
| Supranuclear gaze palsy | 41.5 (27) | 87.7 (57) | 63.6 (21) | 97.0 (32) | 18.8 (6) | 78.1 (25) | **<0.001** | 0.024 |
| Frontal personality change | 36.9 (24) | 69.2 (45) | 39.4 (13) | 66.7 (22) | 34.4 (11) | 71.9 (23) | n.s. | n.s. |
| Non-specific visual symptoms | 35.4 (23) | 67.7 (44) | 42.4 (14) | 69.7 (23) | 28.1 (9) | 65.6 (21) | n.s. | n.s. |
| Postural instability | 32.3 (21) | 86.2 (56) | 42.4 (14) | 93.9 (31) | 21.9 (7) | 78.1 (25) | n.s. | n.s. |
| Frontal physical signs | 27.7 (18) | 80.0 (52) | 24.2 (8) | 78.8 (26) | 31.3 (10) | 81.3 (26) | n.s. | n.s. |
| Dysphagia | 24.6 (16) | 84.6 (55) | 30.3 (10) | 93.9 (31) | 18.8 (6) | 75.0 (24) | n.s. | 0.037 |
| Frontal social dysfunction | 23.1 (15) | 38.5 (25) | 21.2 (7) | 27.3 (9) | 25.0 (8) | 50.0 (16) | n.s. | n.s. |
| Rapid micrographia | 21.5 (14) | 23.1 (15) | 24.2 (8) | 24.2 (8) | 18.8 (6) | 21.9 (7) | n.s. | n.s. |
| Limbs rigidity > axial rigidity | 20.0 (13) | 27.7 (18) | 15.2 (5) | 18.2 (6) | 25.0 (8) | 37.5 (12) | n.s. | n.s. |
| Axial rigidity > limbs rigidity | 18.5 (12) | 55.4 (36) | 33.3 (11) | 69.7 (23) | 3.1 (1) | 40.6 (13) | 0.002 | 0.017 |
| Frontal behaviour | 16.9 (11) | 50.8 (33) | 15.2 (5) | 45.5 (15) | 18.8 (6) | 56.3 (18) | n.s. | n.s. |
| Apraxia of limb(s) | 13.8 (9) | 29.2 (19) | 6.1 (2) | 18.2 (6) | 21.9 (7) | 40.6 (13) | n.s. | 0.042 |
| Progressive non-fluent aphasia | 13.8 (9) | 13.8 (9) | 0 (0) | 0 (0) | 28.1 (9) | 28.1 (9) | **0.001** | **0.001** |
| Pyramidal tract signs | 12.3 (8) | 35.4 (23) | 9.1 (3) | 27.3 (9) | 15.6 (5) | 43.8 (14) | n.s. | n.s. |
| AD-like deficits | 10.8 (7) | 46.2 (30) | 9.1 (3) | 42.4 (14) | 12.5 (4) | 50.0 (16) | n.s. | n.s. |
| Tremor | 10.8 (7) | 21.5 (14) | 18.2 (6) | 30.3 (10) | 3.1 (1) | 12.5 (4) | n.s. | n.s. |
| Limb(s) dystonia | 9.2 (6) | 26.2 (17) | 3.0 (1) | 15.2 (5) | 15.6 (5) | 37.5 (12) | n.s. | 0.038 |
| Rapid hypophonia | 9.2 (6) | 13.8 (9) | 12.1 (4) | 15.2 (5) | 6.3 (2) | 12.5 (4) | n.s. | n.s. |
| Autonomic dysfunction | 7.7 (5) | 46.2 (30) | 6.1 (2) | 42.4 (14) | 9.4 (3) | 50.0 (16) | n.s. | n.s. |
| Cortical sensory loss | 7.7 (5) | 18.5 (12) | 3.0 (1) | 9.1 (3) | 12.5 (4) | 28.1 (9) | n.s. | 0.048 |
| Rest tremor | 6.2 (4) | 9.2 (6) | 12.1 (4) | 18.2 (6) | 0 (0) | 0 (0) | n.s. | 0.013 |
| Freezing of speech | 6.2 (4) | 9.2 (6) | 6.1 (2) | 9.1 (3) | 6.3 (2) | 9.4 (3) | n.s. | n.s. |
| Freezing of gait | 6.2 (4) | 7.7 (5) | 9.1 (3) | 9.1 (3) | 3.1 (1) | 6.3 (2) | n.s. | n.s. |
| Start hesitation | 4.6 (3) | 4.6 (3) | 3.0 (1) | 3.0 (1) | 6.3 (2) | 6.3 (2) | n.s. | n.s. |
| Axial dystonia | 3.1 (2) | 27.7 (18) | 3.0 (1) | 33.3 (11) | 3.1 (1) | 21.9 (7) | n.s. | n.s. |
| Lid apraxia | 3.1 (2) | 20.0 (13) | 0 (0) | 9.1 (3) | 6.3 (2) | 31.3 (10) | n.s. | 0.026 |
| Alien limb phenomenon | 3.1 (2) | 6.2 (4) | 0 (0) | 3.0 (1) | 6.3 (2) | 9.4 (3) | n.s. | n.s. |
| Cerebellar signs | 3.1 (2) | 6.2 (4) | 3.0 (1) | 6.1 (2) | 3.1 (1) | 6.3 (2) | n.s. | n.s. |
| Spontaneous hallucinations | 3.1 (2) | 6.2 (4) | 6.1 (2) | 6.1 (2) | 0 (0) | 6.3 (2) | n.s. | n.s. |
| Myoclonus | 1.5 (1) | 13.8 (9) | 0 (0) | 3.0 (1) | 3.1 (1) | 25.0 (8) | n.s. | 0.012 |
| Apraxia of speech | 1.5 (1) | 4.6 (3) | 0 (0) | 3.0 (1) | 3.1 (1) | 6.3 (2) | n.s. | n.s. |
| Semantic dementia | 0 (0) | 0 (0) | 0 (0) | 0 (0) | 0 (0) | 0 (0) | n.s. | n.s. |
| Unilateral onset | 13.8 (9) |  | 6.1 (2) |  | 21.9 (7) |  | n.s. |  |

|  |  |  |  |  |
| --- | --- | --- | --- | --- |
| Asymmetry at onset | 30.8 (20) | 39.4 (13) | 21.9 (7) | n.s. |
| Persistent asymmetry | 62.1 (18) | 46.7 (7) | 78.6 (11) | n.s. |
| Levodopa responsiveness | 24.3 (9) | 29.2 (7) | 15.4 (2) | n.s. |
| Levodopa-induced dyskinesia | 2.7 (1) | 0 (0) | 7.7 (1) | n.s. |
| Levodopa-induced hallucinations | 0 (0) | 0 (0) | 0 (0) | n.s. |
| Amantadine-induced hallucinations | 37.0 (10) | 41.2 (7) | 30.0 (3) | n.s. |

Fisher's exact test. p refers to the comparison between PSP-RS and vPSP cases at first and last visits. Features are ordered according to decreasing prevalence in the whole PSP cohort, and reported in percentages, with numbers in brackets. p values in bold indicate survival to the multiple comparisons-corrected threshold ( $p < 0.0013$ ).
vPSP: variant PSP syndromes; n.s.: not significant.

**Supplementary Table 5.**
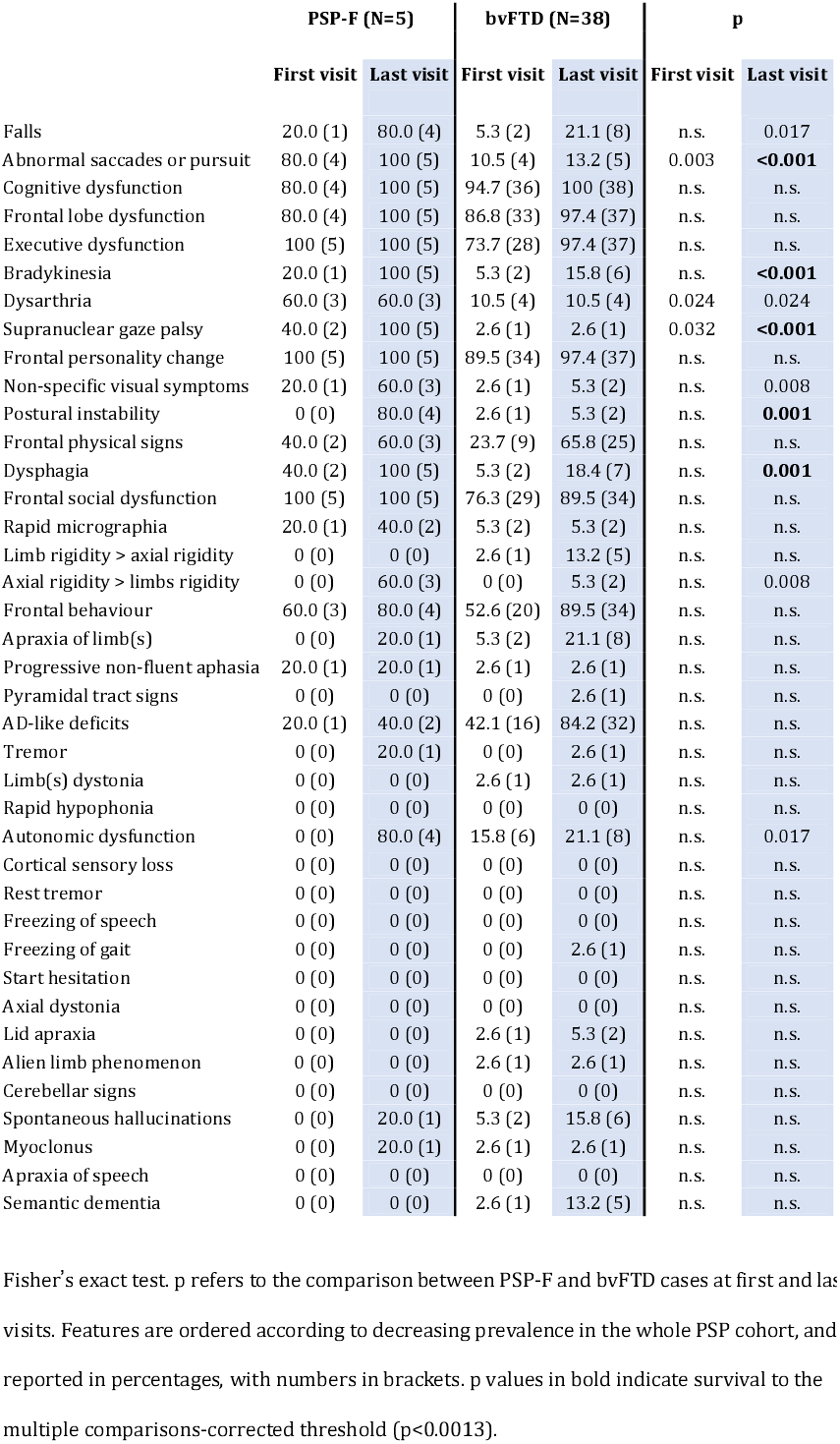
Prevalence of PSP features in PSP-F and bvFTD cases.

**Supplementary Table 6.**
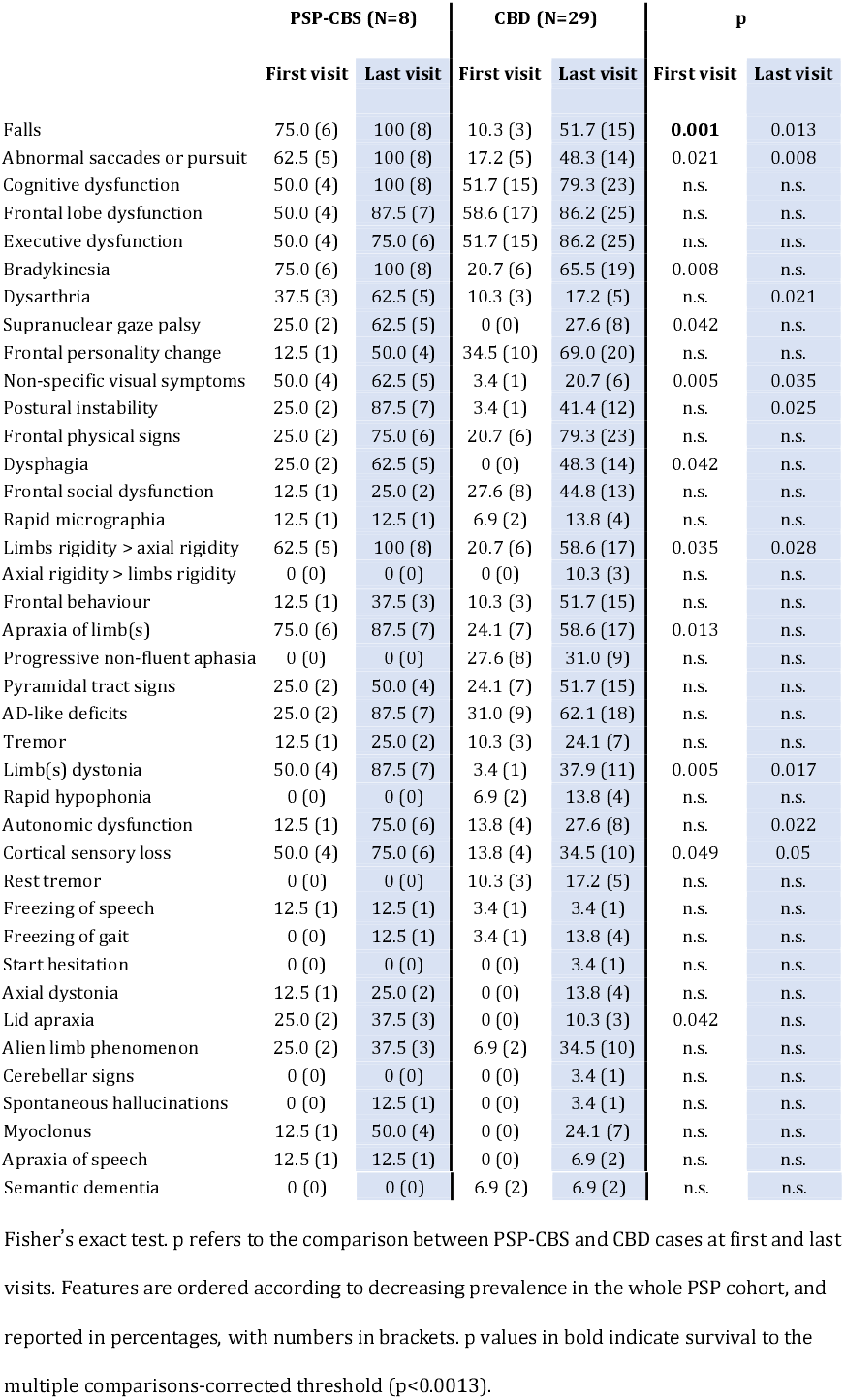
Prevalence of PSP features in PSP-CBS and CBD cases.

**Supplementary Table 7.**
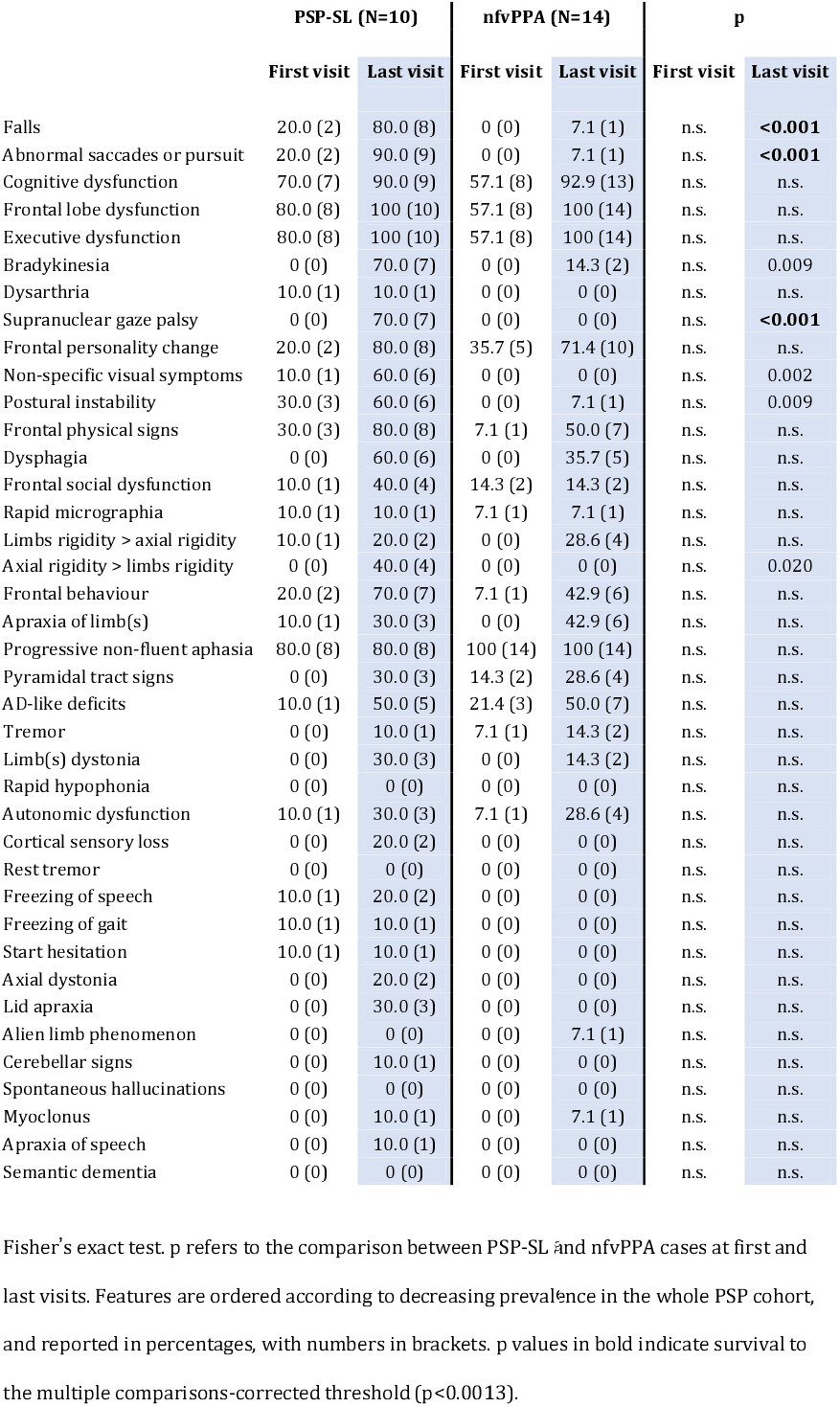
Prevalence of PSP features in PSP-SL and nfvPPA cases.

## BIBLIOGRAPHY

1. Steele JC, Richardson JC, Olszewski J. Progressive Supranuclear Palsy. A Heterogeneous Degeneration Involving the Brain Stem, Basal Ganglia and Cerebellum with Vertical Gaze and Pseudobulbar Palsy, Nuchal Dystonia and Dementia. Arch Neurol 1964;10:333–359.

2. Kovacs GG. Invited review: Neuropathology of tauopathies: principles and practice. Neuropathol Appl Neurobiol 2015;41(1):3–23.

3. Dickson DW. Neuropathologic differentiation of progressive supranuclear palsy and corticobasal degeneration. J Neurol 1999;246 Suppl 2:116–15.

4. Litvan I, Agid Y, Calne D, et al. Clinical research criteria for the diagnosis of progressive supranuclear palsy (Steele-Richardson-Olszewski syndrome): report of the NINDS-SPSP international workshop. Neurology 1996;47(1):1–9.

5. Williams DR, de Silva R, Paviour DC, et al. Characteristics of two distinct clinical phenotypes in pathologically proven progressive supranuclear palsy: Richardson’s syndrome and PSP-parkinsonism. Brain 2005;128(Pt 6):1247–1258.

6. Williams DR, Holton JL, Strand K, Revesz T, Lees AJ. Pure akinesia with gait freezing: a third clinical phenotype of progressive supranuclear palsy. Mov Disord 2007;22(15):2235–2241.

7. Tsuboi Y, Josephs KA, Boeve BF, et al. Increased tau burden in the cortices of progressive supranuclear palsy presenting with corticobasal syndrome. Mov Disord 2005;20(8):982–988.

8. Hassan A, Parisi JE, Josephs KA. Autopsy-proven progressive supranuclear palsy presenting as behavioral variant frontotemporal dementia. Neurocase 2012;18(6):478–488.

9. Boeve B, Dickson D, Duffy J, Bartleson J, Trenerry M, Petersen R. Progressive nonfluent aphasia and subsequent aphasic dementia associated with atypical progressive supranuclear palsy pathology. Eur Neurol 2003;49(2):72–78.

10. Kanazawa M, Shimohata T, Toyoshima Y, et al. Cerebellar involvement in progressive supranuclear palsy: A clinicopathological study. Mov Disord 2009;24(9):1312–1318.

11. Respondek G, Stamelou M, Kurz C, et al. The phenotypic spectrum of progressive supranuclear palsy: a retrospective multicenter study of 100 definite cases. Mov Disord 2014;29(14): 1758–1766.

12. Osaki Y, Ben-Shlomo Y, Lees AJ, et al. Accuracy of clinical diagnosis of progressive supranuclear palsy. Mov Disord 2004;19(2):181–189.

13. Sakamoto R, Tsuchiya K, Mimura M. Clinical heterogeneity in progressive supranuclear palsy: problems of clinical diagnostic criteria of NINDS-SPSP in a retrospective study of seven Japanese autopsy cases. Neuropathology 2010;30(1):24–35.

14. Kanazawa M, Tada M, Onodera O, Takahashi H, Nishizawa M, Shimohata T. Early clinical features of patients with progressive supranuclear palsy with predominant cerebellar ataxia. Parkinsonism Relat Disord 2013;19(12):1149–1151.

15. Hoglinger GU, Respondek G, Stamelou M, et al. Clinical diagnosis of progressive supranuclear palsy: The movement disorder society criteria. Mov Disord 2017;32(6):853–864.

16. Boxer AL, Yu JT, Golbe LI, Litvan I, Lang AE, Hoglinger GU. Advances in progressive supranuclear palsy: new diagnostic criteria, biomarkers, and therapeutic approaches. Lancet Neurol 2017;16(7):552–563.

17. Respondek G, Kurz C, Arzberger T, et al. Which ante mortem clinical features predict progressive supranuclear palsy pathology? Mov Disord 2017;32(7):995–1005.

18. Hauw JJ, Daniel SE, Dickson D, et al. Preliminary NINDS neuropathologic criteria for Steele-Richardson-Olszewski syndrome (progressive supranuclear palsy). Neurology 1994;44(11):2015–2019.

19. Cairns NJ, Bigio EH, Mackenzie IR, et al. Neuropathologic diagnostic and nosologic criteria for frontotemporal lobar degeneration: consensus of the Consortium for Frontotemporal Lobar Degeneration. Acta Neuropathol 2007;114(1):5–22.

20. Dickson DW, Bergeron C, Chin SS, et al. Office of Rare Diseases neuropathologic criteria for corticobasal degeneration. J Neuropathol Exp Neurol 2002;61(11):935–946.

21. Mackenzie IR, Neumann M, Bigio EH, et al. Nomenclature and nosology for neuropathologic subtypes of frontotemporal lobar degeneration: an update. Acta Neuropathol 2010;119(1):1–4.

22. Alexander SK, Rittman T, Xuereb JH, Bak TH, Hodges JR, Rowe JB. Validation of the new consensus criteria for the diagnosis of corticobasal degeneration. J Neurol Neurosurg Psychiatry 2014;85(8):925–929.

23. Gerstenecker A, Mast B, Duff K, Ferman TJ, Litvan I, Group E-PS. Executive dysfunction is the primary cognitive impairment in progressive supranuclear palsy. Arch Clin Neuropsychol 2013;28(2):104–113.

24. Massey LA, Micallef C, Paviour DC, et al. Conventional magnetic resonance imaging in confirmed progressive supranuclear palsy and multiple system atrophy. Mov Disord 2012;27(14): 1754–1762.

25. Saeed U, Compagnone J, Aviv RI, et al. Imaging biomarkers in Parkinson’s disease and Parkinsonian syndromes: current and emerging concepts. Transl Neurodegener 2017;6:8.

26. Bacioglu M, Maia LF, Preische O, et al. Neurofilament Light Chain in Blood and CSF as Marker of Disease Progression in Mouse Models and in Neurodegenerative Diseases. Neuron 2016;91(1):56–66.

27. Magdalinou NK, Paterson RW, Schott JM, et al. A panel of nine cerebrospinal fluid biomarkers may identify patients with atypical parkinsonian syndromes. J Neurol Neurosurg Psychiatry 2015;86(11):1240–1247.

28. Stamelou M, Diehl-Schmid J, Hapfelmeier A, et al. The frontal assessment battery is not useful to discriminate progressive supranuclear palsy from frontotemporal dementias. Parkinsonism Relat Disord 2015;21(10):1264–1268.

29. Santos-Santos MA, Mandelli ML, Binney RJ, et al. Features of Patients With Nonfluent/Agrammatic Primary Progressive Aphasia With Underlying Progressive Supranuclear Palsy Pathology or Corticobasal Degeneration. JAMA Neurol 2016;73(6):733–742.

30. Glasmacher SA, Leigh PN, Saha RA. Predictors of survival in progressive supranuclear palsy and multiple system atrophy: a systematic review and meta-analysis. J Neurol Neurosurg Psychiatry 2017;88(5):402–411.

31. Litvan I, Mangone CA, McKee A, et al. Natural history of progressive supranuclear palsy (Steele-Richardson-Olszewski syndrome) and clinical predictors of survival: a clinicopathological study. J Neurol Neurosurg Psychiatry 1996;60(6):615–620.

32. Nath U, Ben-Shlomo Y, Thomson RG, Lees AJ, Burn DJ. Clinical features and natural history of progressive supranuclear palsy: a clinical cohort study. Neurology 2003;60(6):910–916.

33. Scaravilli T, Tolosa E, Ferrer I. Progressive supranuclear palsy and corticobasal degeneration: lumping versus splitting. Mov Disord 2005;20 Suppl 12:S21–28.

34. Houlden H, Baker M, Morris HR, et al. Corticobasal degeneration and progressive supranuclear palsy share a common tau haplotype. Neurology 2001;56(12):1702–1706.

35. Josephs KA, Petersen RC, Knopman DS, et al. Clinicopathologic analysis of frontotemporal and corticobasal degenerations and PSP. Neurology 2006;66(1):41–48.

